# Genome-wide association and functional analyses identify CASC20 and KIF26B as target loci in heterotopic ossification

**DOI:** 10.1101/845958

**Authors:** Konstantinos Hatzikotoulas, George AE Pickering, Matthew J Clark, Favour Felix-Ilemhenbhio, Klaudia Kocsy, Jonathan Simpson, Scott J MacInnes, Mine Koprulu, Lorraine Southam, Ilaria Bellantuono, Kajus Baidžajevas, David A Young, Alison Gartland, Eleftheria Zeggini, Endre Kiss-Toth, J Mark Wilkinson

## Abstract

Heterotopic ossification (HO) is bone formation that occurs after trauma within tissues that do not normally have the property of ossification, resulting in pain and disability. The genetic architecture of HO remains unclear. In the first genome-wide association studies of this disease, we identify the human-only long non-coding RNA-encoding gene *CASC20* as a robust, replicating susceptibility locus for HO and *KIF26B* as a potential severity locus. We find that both *CASC20* and *KIF26B* are expressed in human bone. Both *CASC20* and *KIF26B* expression is upregulated upon BMP2 induced osteogenic differentiation in primary human mesenchymal stem cells, followed by *RUNX2* and *OSTERIX* upregulation and mineralised nodule formation. A CRISPR-Cas9 mediated knockout of *Kif26b* inhibits BMP2-induced *Runx2*, *Sp7/Osterix*, *Col1A1*, *Alp*, and *Bglap*/*Osteocalcin* expression in a murine myocyte model of osteogenic trans-differentiation, and prevents mineralised nodule formation. These studies provide the first insights into the heritable biology of common, complex HO.

## INTRODUCTION

Heterotopic ossification (HO) is the pathological formation of bone within extra-skeletal tissues that do not normally ossify. HO is a common sequel of trauma, developing in approximately two thirds of casualties after blast injury.^1,2^ It is also common after hip replacement surgery (total hip arthroplasty, THA) with a reported incidence between 5% for more severe disease and up to 90% for milder forms.^3,4^ HO also occurs after traumatic brain injury and burns.^5,6^ The clinical impacts of HO include pain and restricted movement, and symptoms due to the compression of adjacent structures, such as nerves and blood vessels. Clinical risk factors for HO after THA include male sex, hypertrophic osteoarthritis, ankylosing spondylitis, hip ankylosis, and African-American ethnicity that suggest a common, complex aetiology.^7,8^ Currently, there are no effective, mechanism-based treatments for HO, and thus identification of focussed, actionable drug targets is required.

Fibrodysplasia ossificans progressiva (FOP) is a rare but devastating and invariably fatal monogenic disease that is also characterised by bone formation at extra-skeletal sites. In 97% of cases it is caused by a constitutively-activating mutation in the Activin receptor A type I gene (*ACVR1*) that codes for a type I bone morphogenetic protein (BMP) receptor.^9,10^ In contrast, the molecular pathogenesis of common, complex HO is poorly understood, although BMP-induced signalling is also thought to play a role.^11,12^ BMPs are members of the transforming growth factor beta (TGF-ß) superfamily and signal via type 1 and type 2 BMP cell surface receptors.^13–15^ BMP2 is overexpressed in clinically evolving HO tissue after trauma,^16,17^ and BMP type 1 or 2 receptor inhibition reduces HO formation in experimental models.^18,19^ This pathway plays a central role in bone regeneration and repair, inducing the proliferation and differentiation of condensing mesenchymal stem cells (MSCs) towards chondrocytes and osteoblasts. These signals are transduced via mitogen-activated protein kinase (MAPK) signalling^20^ and/or SMAD complexes.^21^ The canonical *Wnt*/ß-catenin pathway is also implicated in chondrocyte maturation and osteoblast differentiation.^22,23^ Both the BMP and *Wnt* pathways converge on the transcription factors *Runx2* and *Sp7* (*Osterix)*, resulting in HO formation through either endochondral or intramembranous routes,^22,24,25^ depending upon the local tissue environment.

Here, we report the first genome-wide association analyses for HO susceptibility and severity in patients after THA. Prioritised signals were followed up in an independent patient cohort. Expression of implicated genes was confirmed in human bone and mechanistically explored using *in vitro* models of osseous transdifferentiation. We found that variation in the long non-coding RNA-encoding gene *CASC20* is robustly associated with HO susceptibility. We also found a putative association between variation within *KIF26B*, a kinesin gene, and the severity of HO development. However, we identified no association signals at or around the *ACVR1* locus. We found that both *CASC20* and *KIF26B* are expressed in human bone, and that both are induced in BMP2-stimulated primary human MSCs, resulting in osseous differentiation. In a murine model of osseous trans-differentiation we found that knockout of *Kif26b* abrogates BMP2-induced bone formation, *Runx2* and *Osterix* expression and synthesis of bone matrix proteins. These effects were associated with dysregulated ERK1/2 MAP-kinase signalling. The identification of susceptibility loci and putative mechanisms in common, complex HO provide a novel opportunity for the development of targeted interventions.

## RESULTS

### Variation within the CASC20 locus is associated with HO susceptibility

To examine the genetic architecture of HO, we conducted genome-wide association analyses for disease susceptibility (cases and controls) and disease severity (within cases) in patients not less than 1 year after total hip replacement for osteoarthritis. Genome wide analysis of the discovery cohort identified an excess of signals associated with HO susceptibility (Figure 1A & 1B, and Supplementary Table 1). Following linkage disequilibrium (LD) pruning using the clumping function in PLINK^26^, we identified 20 independent signals at *p* ≤ 2.68×10^−5^(Supplementary Table 2), thirteen of which passed our variant-level quality control (QC) pipeline for *de novo* replication (see Methods).

**Figure 1.**
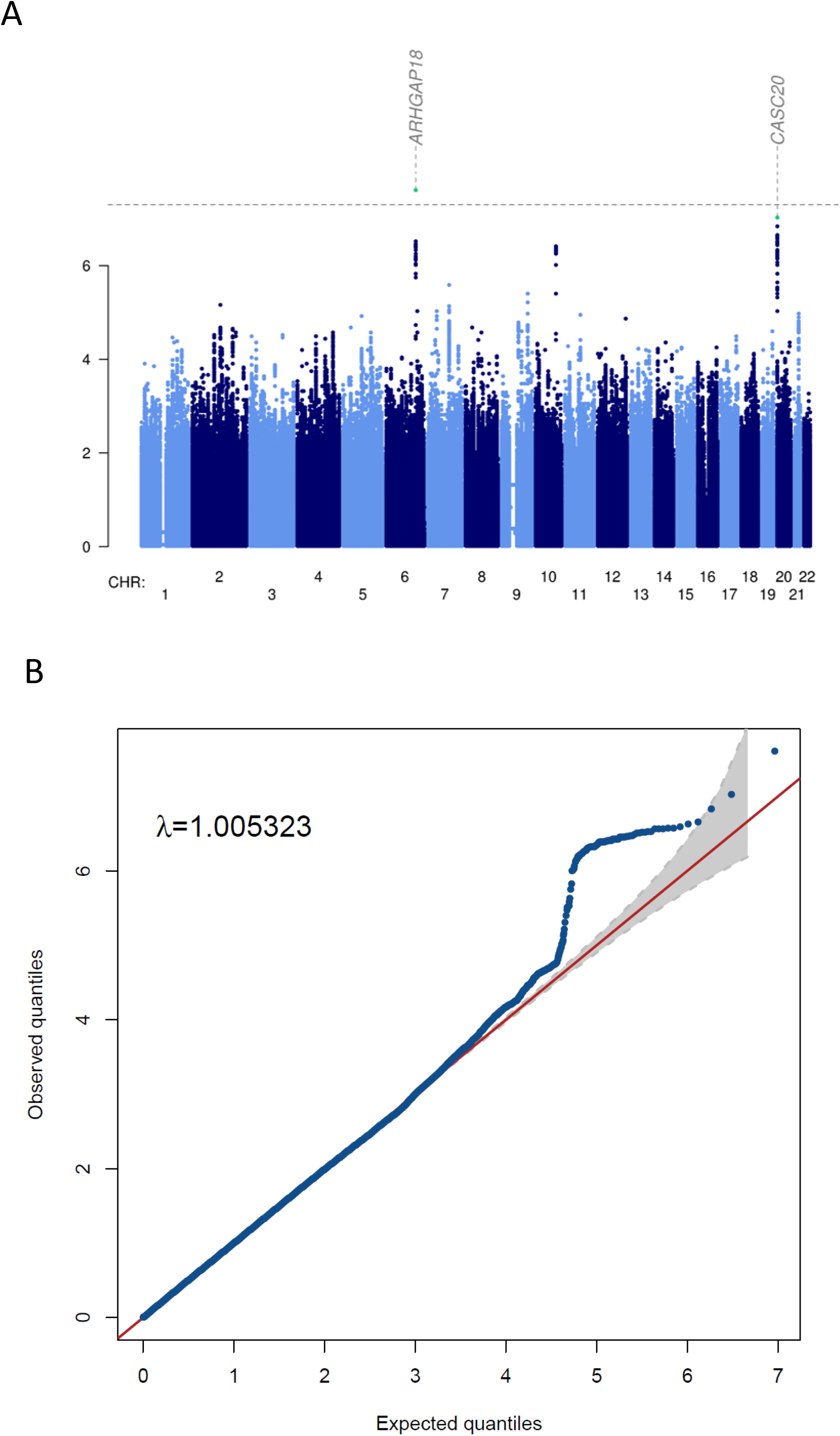
Manhattan and quantile–quantile plot of the discovery HO susceptibility genome-wide association scan. **A)** Manhattan plot showing the −log10 *p*-values for each variant (y axis) plotted against their respective chromosomal position (x axis). The horizontal dashed line denotes the genome-wide significance threshold *p*=5×10^−8^. **B)** QQ plot where the x-axis indicates the expected −log10 *p*-values and the y-axis the observed ones. The red line represents the null hypothesis of no association at any locus and λ is the genomic inflation factor.

The strongest signal, which reached genome wide significance, lay in an intergenic region just downstream of *ARHGAP18* that encodes a protein involved in the modulation of cell signalling, cell shape and motility.^27^(rs59084763, effect allele (EA) T, effect allele frequency (EAF) 0.19, OR [95% CI] 1.87 [1.47–2.37], *p*=2.48×10^−8^; Figure 2A). The second strongest signal lay within the long non-coding (LNC) RNA-encoding gene *CASC20* (rs11699612, EA T, EAF 0.25, OR 1.73 [1.40-2.16], *p*=9.39×10^−8^; Supplementary Figure 1). *CASC20* is a human only LNC RNA that has no orthologues in species outside apes. Both its mechanism of action and functional importance in health and disease are currently unexplored. Another strong signal, approaching genome wide significance (rs10882328, EA A, MAF 0.29, OR 0.58 [0.47-0.71], *p*=3.87×10^−7^; Supplementary Figure 2) lay within *LGI1* that encodes the Leucine-rich, glioma inactivated 1 protein, and is expressed in neural tissue.^28^

**Figure 2.**
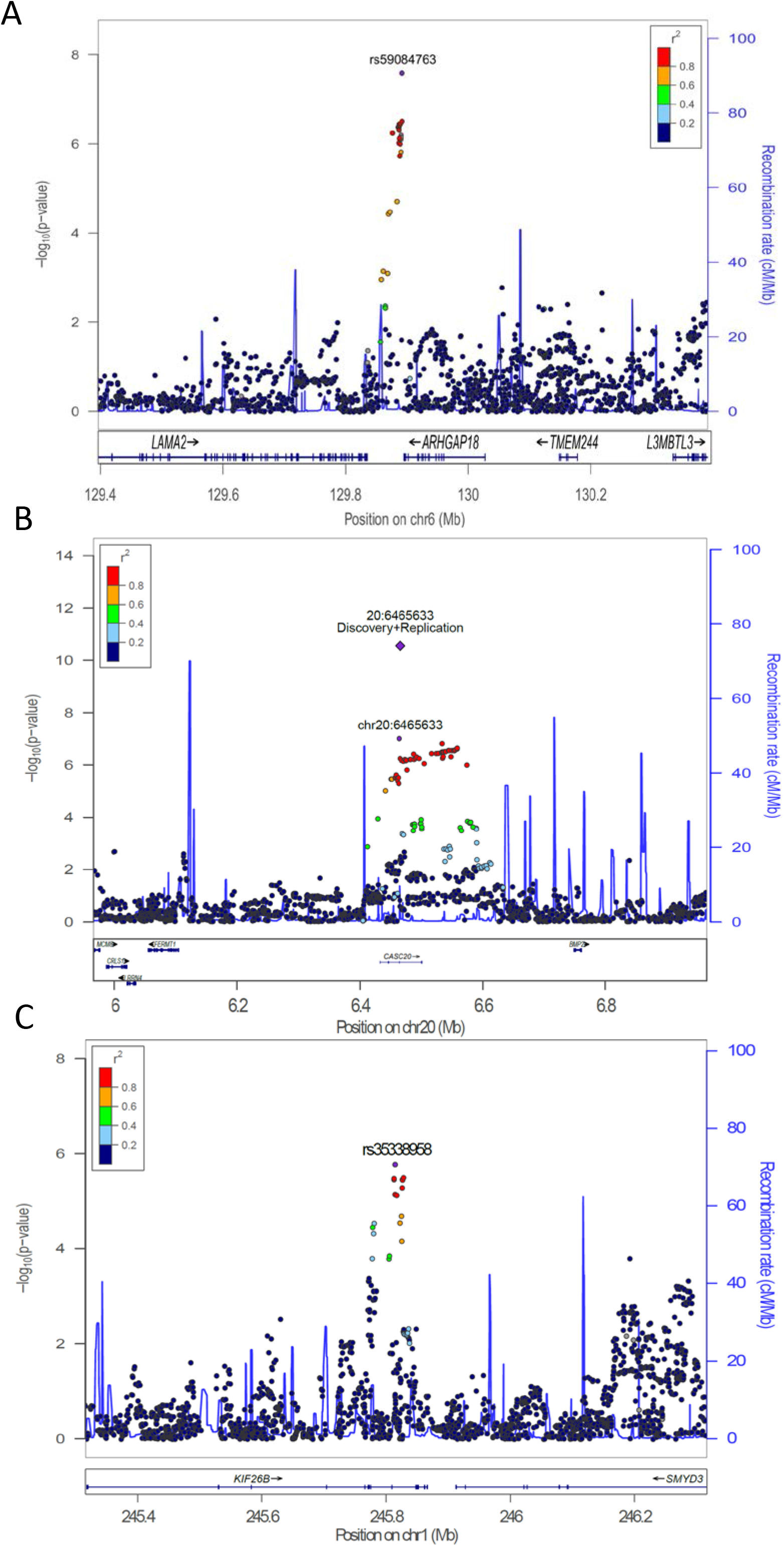
Association of lead signals with heterotopic ossification. A) Regional association plot of rs59084763 with HO susceptibility. Each filled circle represents the *p*-value of analysed variants in the discovery stage plotted against their physical position (NCBI Build 37). The purple circle denotes rs59084763 which is the variant with the lowest *p*-value in the region. **B) Regional association plot of the CASC20 variant with HO susceptibility.** Each filled circle represents the *p*-value of analysed variants in the discovery stage plotted against their physical position (NCBI Build 37). The *p*-value at the discovery stage and combined discovery and replication cohorts of rs11699612 is represented by a purple circle and diamond, respectively. **C) Regional association plot of the KIF26B variant association with HO severity.** Each filled circle represents the *p*-value of analysed variants in the discovery stage plotted against their physical position (NCBI Build 37). The purple circle denotes rs35338958 which is the variant with the lowest *p*-value in the region. The colours of variants in each plot indicate their r^2^ with the lead variant according to a scale from r^2^ = 0 (blue) to r^2^ = 1 (red).

At replication in an independent THA patient cohort, four of the thirteen independent HO susceptibility signals that passed variant-level QC (Supplementary Table 2) showed a concordant direction of effect and one residing within the *CASC20* locus replicated at the corrected significance threshold (rs11699612, EA T, EAF 0.23, OR 1.90 [1.34-2.68], *p*=2.70×10^−5^; Supplementary Table 3). Following meta-analysis, this association reached genome-wide significance (EA T, EAF 0.24, OR 1.94 [1.59-2.35], *p*=2.71×10^−11^; Figure 2B). *CASC20* resides in close proximity to *BMP2* on chr20. In order to confirm the origin of the signal as lying within *CASC20* rather than *BMP2*, we increased the physical distance threshold used for clumping to 2000 kb either side of rs11699612 to include the whole *BMP2* locus and most of the proximal coding genes. We found no HO-associating variants within *BMP2* nor any *BMP2* variants in higher LD than r^2^=0.05 with the *CASC20* variant, confirming *CASC20* as the origin of the signal (Supplementary Table 4). Given the proximity of *CASC20* to *BMP2*, we further examined whether rs11699612 is a cis-acting expression quantitative trait locus (cis-eQTL) for *BMP2* expression using RNA sequencing data from unstimulated primary chondrocytes and synoviocytes taken from an independent cohort of 100 patients undergoing joint replacement (See data availability section and bioRxiv manuscript biorxiv.org/content/10.1101/835850v1). Data were analysed using a GTEx modified version of FastQTL,29 (PMID: 26708335) and confirmed no evidence of rs11699612 action as a cis-eQTL on *BMP2* in chondrocytes, nor evidence of any other cis-acting eQTLs in *BMP2* (Supplementary Table 5).

### Variation within KIF26B and disease severity

Twenty-two signals were suggestively associated with HO severity, with clumped index variants at *P*<2.68×10^−5^, the most significant 10 of which were prioritised for replication (Supplementary Tables 6 and 7). The strongest signal lay within an intronic region of *KIF26B* (rs35338958, EA T, EAF 0.10, OR 3.04 [1.85-5.01], *p*=1.65×10^−6^; Figure 2C). Whilst no signals in the severity association analysis reached either the corrected *p*-value threshold in the replication stage or genome-wide significance threshold in the meta-analysis, eight out of the selected ten independent signals showed a concordant direction of effect between the discovery and replication stages, including the intronic *KIF26B* variant rs35338958. Given previous literature suggesting a mechanistic role for *KIF26B* in ectopic calcification^30^ and in WNT signalling^31^, we next explored the role of both *CASC20* and *KIF26B* as candidate loci in subsequent functional analyses.

### CASC20 and KIF26B are expressed in human bone, are induced in mammalian HO models, including in BMP2 stimulated human mesenchymal stem cells

To determine whether our lead susceptibility and severity genes are expressed in human bone tissue, we extracted total RNA from fresh frozen, surgically excised bone from patients undergoing joint replacement and confirmed expression of both *CASC20* and *KIF26B* expression by real time quantitative polymerase chain reaction (RT-qPCR, Figure 3A). Next, we explored whether *KIF26B* and *CASC20* are differentially expressed in human multipotent adipose-derived stem cells (hMAD, Figure 3B)^32^ and in primary human bone marrow-derived mesenchymal stem cells (hMSCs, Figure 3C-E) in response to stimulation with BMP2 and osteogenic supplements. Runt-related transcription factor 2 (*RUNX2*) and the transcription factor Sp7/Osterix (*OSX*), markers of osteogenic differentiation, were robustly upregulated in both the hMADs and the hMSCs. We also identified an increase in percentage mineralisation per well at day 24 measured by Alizarin Red S staining. We observed variation in both magnitude and time-course of *CASC20* and *KIF26B* expression between donors. This was also reflected in the expression of *RUNX2* and *OSX*, although the sample size was insufficient to explore these relationships statistically.

**Figure 3.**
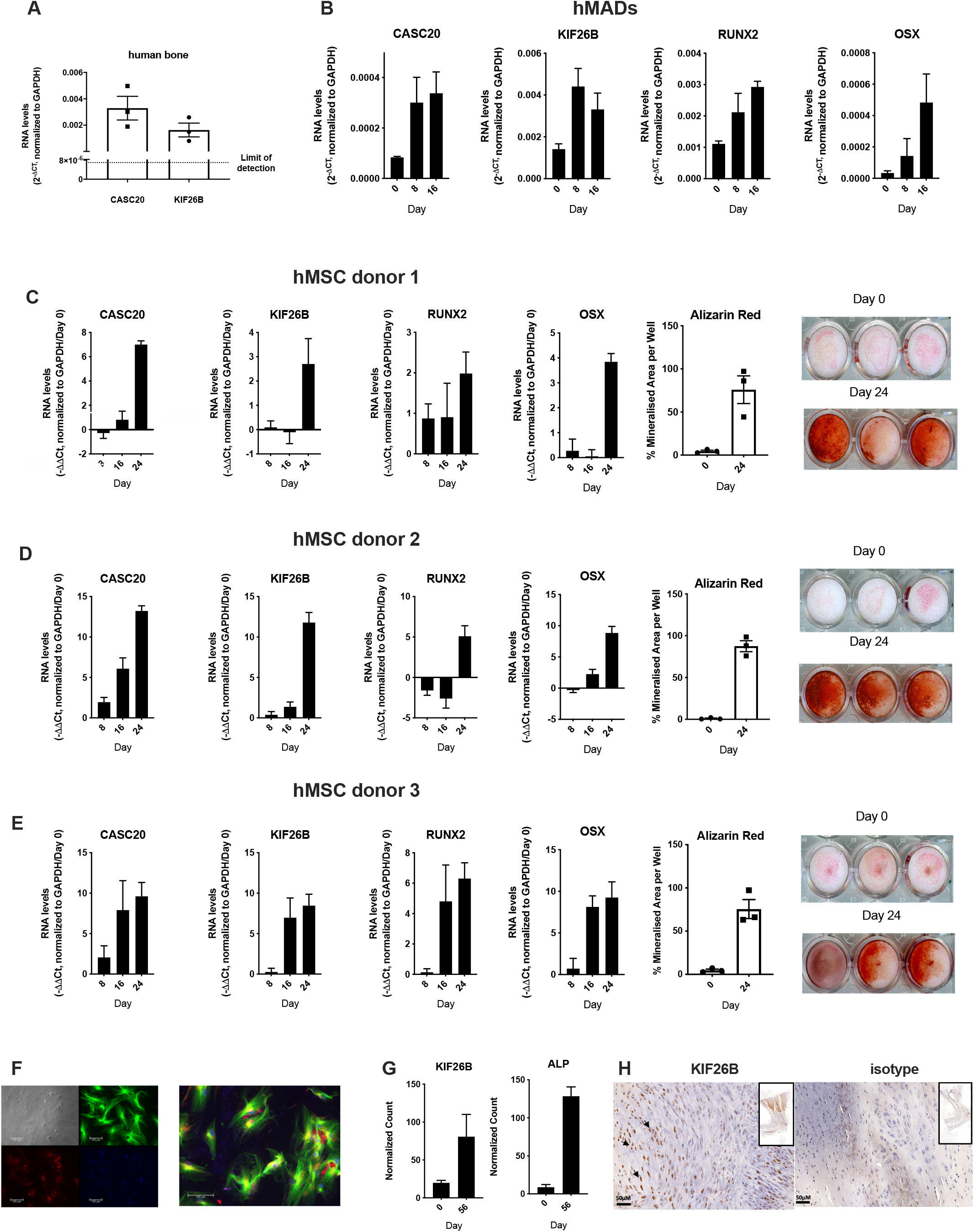
*CASC20* and *KIF26B* are expressed in human bone and are induced in mesenchymal stem cells by BMP2 in vitro and in mammalian models of developing HO tissue. RT-qPCR was used to measure the expression of *CASC20* and *KIF26B* RNA in waste bone samples retrieved at joint replacement (n=3 subjects). **B) *CASC20* and *KIF26B* expression is induced in human Multipotent Adipose-Derived Stem cells (hMADs).** RT-qPCR was used to measure the expression of *CASC20*, *KIF26B, RUNX2* and *OSX* in hMADS at days 0, 8 and 16 of hMAD differentiation. Data were analysed using 2^(-∆Ct) by normalizing to *GAPDH* (n=3 biological replicates). **C-E) CASC20 and KIF26B expression and mineralisation is induced in BMP2-stimulated human Bone Marrow Derived Mesenchymal Stem Cells (hMSC).** RT-qPCR was used to measure the expression of *CASC20*, *KIF26B, RUNX2* and *OSX* in hMSCs from 3 donors at days 0, 8, 16 and 24 of differentiation. Data were analysed using –∆∆Ct, normalizing to day 0 and GAPDH. (n=3 biological replicate cultures) Percentage mineralised area per well at day 0 and day 24, measured by percentage Alizarin Red S staining. Images show Alizarin Red S calcium staining of hMSCs after 0 and 24 days of differentiation. For each panel data are plotted as mean ± SEM. **F) KIF26B protein is expressed in hMSCs under osteogenic conditions.** Immunofluorescence staining of KIF26B (red), Tubulin (green) and DAPI (blue) in hMSCs (from donor 3) differentiated for 8 days. **G) *KIF26B* RNA is upregulated in an *in vivo* model of ectopic bone formation.** Normalised count of *KIF26B* and Alkaline Phosphatase (ALP) compared in days 0 and 56 of differentiation of hMSCs implanted into nude mice (GSE89179). **H) KIF26B protein is expressed in regions of new bone formation in post tendo-achilles injury in rat.** (left) anti-KIF26B antibody staining in regions of new bone formation in the rat Achilles tendon region (right) IgG control antibody staining. Arrows indicate the distinct nuclear localisation of KIF26B (brown) to cells in left panel, including what appears to be hypertrophic chondrocytes. Inset for both panels is a lower magnification of the full section. Scale bar = 50μM.

In addition to upregulation of *KIF26B* and *CASC20* expression and the observed mineralisation, KIF26B protein was also robustly expressed in hMSCs after osteogenic differentiation (Figure 3F). Previous investigators have reported that KIF26B localises to microtubules in human endothelial cells.^33^ Our data confirm that KIF26B has at least a partial co-localisation with the cytoskeleton in MSCs (Figure 3F). *In silico* analysis^34^ of a published RNASeq dataset in an *in vivo* model of ectopic bone formation using human foetal MSCs from an independent study^35^ also demonstrated *KIF26B* induction during osteogenesis (Figure 3G). To study the *in vivo* expression of KIF26B protein following injury, we used immunohistochemistry to identify and co-localise KIF26B expression with HO formation in archived tissue from a rat tendo-Achilles scalpel-induced injury model.^36,37^ In these tissues taken at 10 weeks after injury, islands of new bone formation were identified and co-localised with KIF26B expression (Figure 3H).

### Knockout of murine *Kif26b* prevents BMP2-mediated mineralisation of C2C12 myoblasts

Whilst *in vitro* models that precisely recapitulate the injury and pathogenesis of clinical HO are lacking, C2C12 mouse myoblasts are a commonly used proxy.^38,39^ To substantiate the functional impact of KIF26B during osteogenic trans-differentiation, C2C12 mouse myoblasts were cultured following BMP2 stimulation, leading to a robust increase in *Kif26b* RNA from day 8 (Figure 4A). Of note, *Kif26b* RNA was very low or completely absent in undifferentiated cells at day 0. This absence of expression was further verified at the protein level, as was its upregulation after 8 days of osteogenic differentiation. *Kif26b* RNA was only expressed in the presence of BMP2 plus osteogenic supplements and was not detected with either low serum media, or with osteogenic supplements alone (Figure 4C, wild type cells).

**Figure 4.**
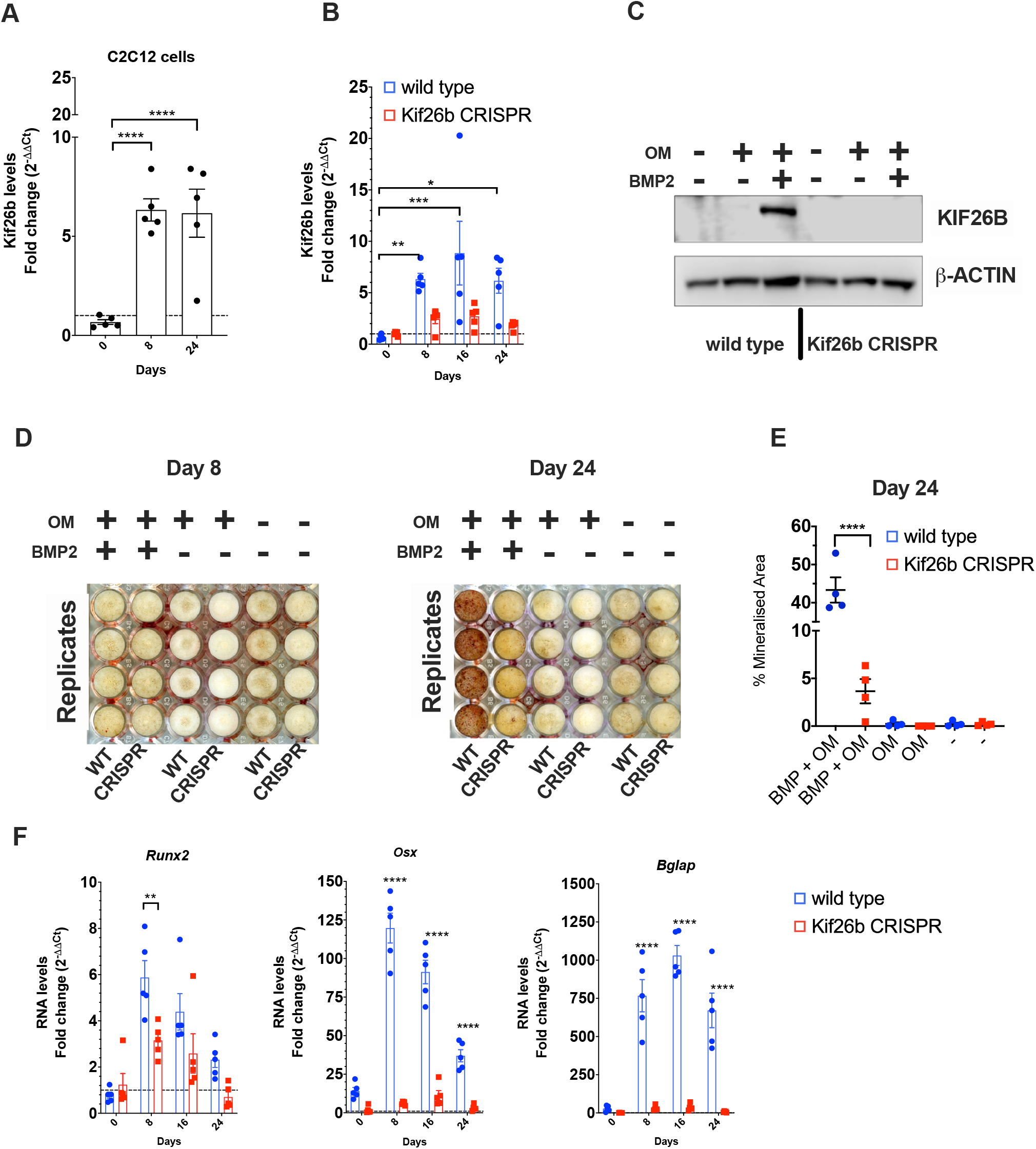
KIF26B is necessary for BMP-induced osteogenic differentiation of C2C12 murine myoblasts. **A) *Kif26b* mRNA expression is induced in C2C12 cells during BMP2 mediated differentiation**. RT-qPCR was used to measure the expression of *Kif26b* at day 0 (confluency) relative to days 8 and 24 of differentiation when treated with 300ng/ml BMP2 and osteogenic supplements. Technical replicates from a representative experiment (from 3 independent differentiations): each data point represents an individual well. One-way ANOVA with Dunnett's multiple comparisons; data is plotted as mean ±SEM. **B) Upregulation of *Kif26b* RNA is abrogated in *Kif26b*^*CRISPR*^ C2C12 cells.** RT-qPCR was used to measure the expression of *Kif26b* at days 8, 16 and 24 of differentiation when treated with 300ng/ml BMP2 and osteogenic supplements relative to day 0 (confluency). Technical replicates from a representative experiment (from 3 independent differentiations): each data point represents an individual well. One-way ANOVA with Dunnett's multiple comparisons; data is plotted as mean ±SEM. **C) Upregulation of KIF26B protein is abrogated in *Kif26b*^*CRISPR*^ C2C12 cells.** KIF26B western blot of WT and *Kif26b*^*CRISPR*^ C2C12 myoblasts that were differentiated for 8 days in three different media. OM: media with osteogenic supplements. **D-E) *In vitro* mineralisation is inhibited in *Kif26b*^*CRISPR*^ C2C12 cells.** D) Alizarin Red S calcium staining of WT and *Kif26b*^*CRISPR*^ C2C12 myoblasts after 8 and 24 days of differentiation. E) Percentage mineralised area per well at day 24, as measured by percentage Alizarin Red S staining. One-way ANOVA with Tukey’s post-hoc test. **F) Induction of osteogenic genes is prevented in *Kif26b*^*CRISPR*^ C2C12 cells.** Cells were cultured in media with osteogenic supplements and BMP2, and RNA levels of *Runx2, Osterix (Osx)* and *Osteocalcin (Bglap)* were measured by qRT-PCR. Two-way ANOVA with Sidak's multiple comparisons. \**P* <0.05, \*\**P* <0.01, \*\*\**P*<0.001, \*\*\*\**P*<0.0001

In cells engineered with CRISPR/Cas9 to not produce KIF26B, there was no expression of *Kif26b* RNA with BMP2 stimulation, in contrast to that observed in the wild-type C2C12 myoblasts (Figure 4B). In the *Kif26b*^*CRISPR*^ cells, mRNA expression was close to the limit of detection at all time-points studied. Western blot confirmed that no KIF26B protein was produced by the *Kif26b*^*CRISPR*^ cells after 8 days of BMP2 stimulation (Figure 4C). Cells of different passage number were stimulated with BMP2 in independent experiments and lysed for western blotting (n=3), confirming the stability of the *Kif26b*^*CRISPR*^ in the C2C12 myoblasts. C2C12 myoblasts express genes promoting osteogenesis and produce mineral when stimulated with BMP2.^38^ Here, we examined the effects of *Kif26b*^*CRISPR*^ on the ability of these cells to produce mineral in a 24-day differentiation experiment using Alizarin Red S staining at days 8 and 24 post-confluency. Figure 4D shows that very little mineral is produced by day 8 in either the WT or *Kif26b*^*CRISPR*^ cells. By day 24, only a small amount of mineral is produced by the *Kif26b*^*CRISPR*^ cells whilst the WT cells produce abundant mineral (3.7% vs 43.3% mean mineralisation, figure 4D-E).

### *Kif26b* knockout disrupts the mineralisation and trans-differentiation of C2C12 myoblasts by modulating the expression of osteogenic genes

Given the lack of mineralisation in the *Kif26b*^*CRISPR*^ cells, we hypothesised that the expression of genes driving osteogenesis would be abrogated. RT-qPCR analysis of *Runx2*, *Osx* and osteocalcin (*Bglap*) at day 0 (confluency), day 8, 16 and 24 was conducted to test this. At day 0, there was no difference in target gene expression between WT and *Kif6b^CRISPR^*. The greatest differential gene expression between cell types was observed at day 8 (Figure 4F). Expression of *Runx2* peaked at day 8 in the WT cells (WT versus *Kif26b*^*CRISPR*^ cells, *p*=0.0052). *Osx* expression in WT cells increased relative to the *Kif26b*^*CRISPR*^ cells at day 8, 16 and 24 (*p*<0.0001 at all treatment timepoints). We also found that Osteocalcin was upregulated in the WT cells but not expressed in *Kif26b*^*CRISPR*^ cells at day 8, 16 and 24 (Figure 4F; *p*<0.0001),

### KIF26B deficiency inhibits ERK MAP kinase activation during osteogenesis, whilst augmenting p38 and SMAD 1/5/8 phosphorylation

In order to understand the molecular mechanism for the abrogated osteogenic response to BMP2 stimulation in *Kif26b*^*CRISPR*^ cells, the activity of key BMP2-effector pathways was tested in wild type and *Kif26b*^*CRISPR*^ cells before and after osteogenic differentiation (Figure 5A-B). The activation of ERK MAPK is critical to osteoblast differentiation in mouse models,^40^*Runx2* and osteocalcin/*Bglap* being major targets of this pathway.^41–43^ In line with these studies, we found that phosphorylation of ERK during osteogenesis was profoundly inhibited in the absence of KIF26B protein (Figure 5A-B). At the same time, phosphorylation of SMAD1/5 and p38 MAPK were not negatively affected in *Kif26b*^*CRISPR*^ cells, suggesting that *Kif26b* is not a critical regulator of these signalling axes in our BMP2-driven model. Mechanistically, RUNX2 has been shown to co-localise with SMADs and form part of a transcriptional activator complex to initiate BMP2 induced osteogenesis.^44^ We thus speculate that the impaired *Runx2* expression seen in *Kif26b*^*CRISPR*^ cells may be sufficient to inhibit osteogenesis. A summary of the proposed functional mechanisms for KIF26B is shown in Figure 5C.

**Figure 5.**
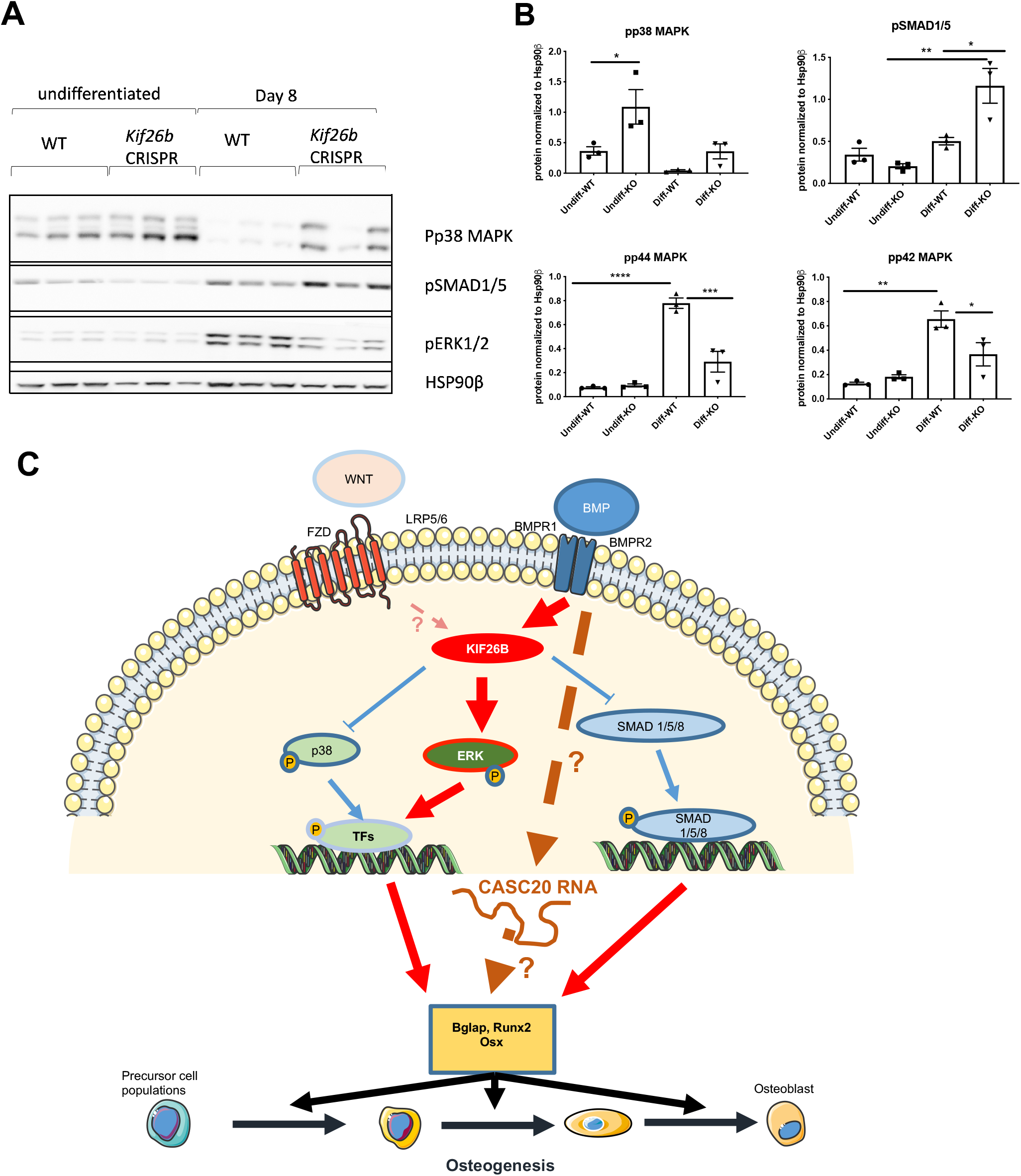
KIF26B-deficiency leads to dysregulated osteogenic signalling, impairing the activation of ERK MAPKs. **A-B)** p38 MAPK, SMAD1/5 and ERK1/2 phosphorylation in WT and *Kif26b*^*CRISPR*^ cells was assessed by western blotting. Cells were differentiated for 8 days (WT/KO) (n=3 biological replicates). B) Western blot data were quantified using Image Studio Lite Ver 5.2. One-way ANOVA with Tukey's multiple comparison test. Data is plotted as mean ±SEM. **C) Proposed model for the molecular mechanism of KIF26B and CASC20 in osteogenesis**. Both KIF26B and CASC20 expression is induced by BMP2. Whilst the mechanistic contribution of CASC20 to HO is unclear, KIF26B appears to control the balance of signalling pathways that collectively regulate then expression of osteogenic genes.

## DISCUSSION

The development of HO is a clinically important sequel of musculoskeletal injury, and for which we currently have only blunt therapeutic tools (radiotherapy or non-steroidal anti-inflammatory agents) that have significant off-target effects. Here, we conducted genome-wide association analyses for HO and identify a robustly replicating locus for HO susceptibility within the previously uncharacterised human-only LNC RNA-encoding gene *CASC20*. We demonstrate its expression in bone and differential expression in human MSC populations in response to an osteogenic stimulus, resulting in mineralised nodule formation. We also find suggestive evidence of association between variance in the kinesin gene *KIF26B* and the severity of formed HO. Based upon this finding, and prior evidence of *Kif26b* expression in an *in vitro* murine model of ectopic ossification^30^ and in WNT signalling,^31,45^ we demonstrate *KIF26B* expression in human bone and its protein in a rat model of HO; and its differential expression in human MSCs in response to an osteogenic stimulus, resulting in mineralised nodule formation. We further show that modulation of KIF26B in a murine myoblast model is sufficient to inhibit osseous trans-differentiation, an effect that is associated with ERK1/2 dysregulation.

p38 MAPK has also been linked to osteogenic differentiation. However, the direction of its effect in this process remains unclear. Whilst a number of studies have shown that p38 activity is required for osteogenesis downstream of BMP2,^20,46^ it has also been reported that inhibition of this pathway can augment osteogenic differentiation.^47^ A recent study showed that TNF inhibited BMP2-mediated osteogenesis via a strong activation of p38 MAPK in C2C12 and MC3T3-E1 cells whilst inhibition of p38 rescued the expression of RUNX2 and other osteogenic markers induced by BMP2.^48^ Our analysis of the consequences of *Kif26b*-deficiency demonstrate over-activity of this pathway, which may also contribute to the observed lack of osteogenesis.

Within the power constraints of our study, we found no evidence to implicate variation within *BMP2* or *ACVR1*, suggesting that the rare disorder FOP does not provide an aetiological cue in common, complex HO. Although both the discovery and replication sample sizes were limited, the genotyped collections studied here amass the largest available sample size globally. Larger sample sizes will be necessary to increase the number of HO risk loci robustly identified, and to discover disease severity signals (Supplementary Figure 3). Further studies of the gene encoding *CASC20* in relevant human cell models will help clarify the role of CASC20 in human health and disease and in the pathogenesis of HO.

We used the C2C12 myoblast cell line to model HO *in vitro*, as muscle precursor cells may contribute to the cellular origin of HO.^49^ This is a sub-clone of a line originally derived from traumatised mouse muscle tissue.^50,51^ This immortalised cell line will readily differentiate into myotubes, adipoblasts or osteoblasts when stimulated under appropriate conditions.^34,47^ When stimulated with BMP2, this cell line is useful in the study of osteogenic differentiation and particularly in the context of HO.^18,39,52–57^ Whilst the mechanistic insight from this cellular model has been valuable, this approach also has limitations. It recapitulates a single cell of origin picture of acquired HO that is known to be complex and involving the interaction of multiple cell types of different embryonal lineages, and the dominant initiating cell type remains unknown.

In summary, here we provide first insights into the genetic architecture of common, complex HO, revealing a susceptibility locus within a LNC RNA encoding gene, and early mechanistic insight into BMP2 modulated signalling in its pathogenesis that may provide opportunities for further investigation as potentially actionable targets in its prevention.

## METHODS

### Study oversight and populations

For the cohort analyses, all subjects were recruited as part of previous ethically approved studies,^58,59^ and provided written, informed consent obtained prior to participation. For the *in vitro* analyses, samples were collected under ethics approval from Oxford NHS REC C (10/H0606/20 and 15/SC/0132), Yorkshire and Humber REC (13/YH/0419), and Human Tissue Authority license 12182, South Yorkshire and North Derbyshire Musculoskeletal Biobank, University of Sheffield, UK.

#### Discovery cohort

891 British Caucasian men and women (481 controls and 410 cases) who had previously undergone hip replacement for idiopathic osteoarthritis were studied.^59^ Controls comprised subjects who had no evidence of HO on plain AP radiographs of the pelvis taken not less than 1 year following primary THA. Cases comprised subjects with radiographic evidence of post-operative HO and were graded (0-4) using the Brooker classification.^60^ Brooker described: a small island of bone (class 1); bone spurs from pelvis and proximal femur leaving at least 1cm between opposing surfaces (class 2); bone spurs from pelvis and proximal femur leaving gap less than 1cm (class 3); apparent ankylosis of the hip (class 4). The distribution of patient demographics and the presence and severity of HO by Brooker classification are shown in Supplementary Table 8.

#### Replication cohort

419 British Caucasian subjects (207 cases and 212 controls) were recruited not less than 1 year following hip replacement for idiopathic osteoarthritis. HO was graded from plain pelvic radiographs as outlined above using the Brooker grading. Cohort demographics and distribution of Brooker grades for the samples proceeding to case control association analyses are shown in Supplementary Table 8.

### Discovery GWAS

DNA from subjects in the discovery cohort was extracted from either whole blood or saliva and genotyped using the Illumina 610k beadchip (Illumina, San Diego, CA). Standard GWAS QC was conducted at the samples and variants level. The exclusion criteria have been previously described.^59,61^ Briefly, individuals with 1) gender discrepancy; 2) call rate <95% (<97% in arcOGEN^61^); 3) excess homozygosity or heterozygosity (more than ±3 SD of the mean); 4) duplicates and related samples (π^>0.2); 5) non-UK European ancestry (ethnicity outliers) were excluded from further analyses. Variants with 1) minor allele frequency (MAF) ≥5% and a call rate <95%, or a MAF <5% and call rate <99%; 2) monomorphic; 3) exact Hardy Weinberg Equilibrium (HWE) p<0.05 (p<0.0001 in arcOGEN) were also excluded from the merged dataset. Variant QC was carried out on autosomal variants. Genotype-calling intensity plots were examined and single nucleotide polymorphisms (SNPs) with poorly clustering plots were not taken forward. Following QC, 891 subjects (481 controls and 410 cases) and 448770 variants were imputed with IMPUTE2^62^ using the European reference panel from the 1000 Genomes Project (Dec 2010 phase I interim release).^63^ Variants with an imputation information score <0.4 and MAF <0.05 were excluded from further analysis.

An HO susceptibility case-control analysis and an evaluation of disease severity using a binomial analysis of Brooker grades 1 and 2 versus grades 3 and 4 (within cases only) were undertaken on >10 million variants under the additive model using method score implemented in SNPTESTv2.^62^ Analyses were adjusted for age and sex as they are known risk factors for HO.^64^ Data were pruned for linkage disequilibrium (LD) using the clumping function in PLINK.^26^ Parameters used: (a) significance threshold for index SNP: 1e-5, (b) LD threshold for clumping: 0.20, and (c) physical distance threshold for clumping: 500 kb. Statistical independence of the signals were also confirmed through conditional single-variant association analyses as implemented in SNPTESTv2. A variant was considered independent of the index SNP if the pre- and post-conditioning *p*-value difference was lower than two orders of magnitude. The first top twenty and ten index SNPs for case-control and severity analyses respectively, were prioritised for replication.

### Replication and meta-analysis

DNA from the subjects in the replication cohort was extracted from saliva and genotyped using the iPLEX® Assay of the MassARRAY® System (Agena Bioscience, Inc) to conduct *de novo* replication. Thirty independent and prioritised variants of the discovery stage (20 susceptibility and 10 severity loci; Supplementary Table 9) were genotyped. All variants had high Agena design metrics and thus were not replaced with highly correlated proxies. QC was conducted at the sample and variant levels. Sample exclusions were based on sex inconsistencies and a sample call rate <60%. Variants with a call rate <75%, minor allele frequency (MAF) <1% and exact HWE *P*<0.001 in controls were also excluded. Following QC, 205 controls and 198 cases and 23 variants (13 susceptibility and 10 severity) proceeded to association analyses. An HO susceptibility case-control analysis and an evaluation of disease severity using a binomial analysis within cases were undertaken under the additive genetic model using the “method maximum likelihood” option as implemented in SNPTESTv2.5.2.^62^ Age and sex were used as covariates.^64^ The significance threshold for association in the replication study was 0.05/23=0.0022. Finally, we performed a fixed-effects inverse-variance-weighted meta-analysis in METAL^65^ across the discovery and replication datasets, comprising a total of 1294 (608 cases and 686 controls) and 608 (136 severe cases and 472 mild cases) individuals for case-control and severity analyses, respectively. Genome-wide significance was defined as *p*<5.0 × 10^−8^.

### Cell culture

Human mesenchymal stem cells (hMSCs) from 3 independent subjects were obtained from the bone marrow of children undergoing osteotomy and cultured in growth medium which consisted of: DMEM containing L-Glutamine (61965-059) and 4.5 g/L Glucose (Gibco), with the addition of 10 % hyclone (SH30070.03). Human multipotent adipose-derived stem cells (hMADS) were cultured in growth medium which consisted of: DMEM (Lonza, BE12-707F), 10% FBS, 1% glutamine (Gibco, 25030-024), 0.2% Penicillin-Streptomycin (P/S), 1% HEPES (Gibco, 15630-056) and 0.01% hFGF2 (Invitrogen, F0291). Confluency was avoided to prevent differentiation, with hMSCs and hMADS cells split every 3-4 days. For osteogenic differentiation studies, cells were cultured in growth medium for 24 hours, followed by 48 hours of exposure to BMP2. At this timepoint the cells were confluent, and the media was changed into differentiation media.

Cells from the C2C12 mouse myoblast cell line (Sigma-Aldrich, St Louis, MO) were cultured in growth medium consisting of: DMEM containing L-Glutamine and 4.5 g/L Glucose (Lonza), with the addition of 10 % Foetal Bovine Serum (FBS), 1 % Penicillin-Streptomycin (P/S) and 1 % L-Glutamine (L-G).

### CRISPR-Cas9 knockout

A CrispR-Cas9 knockout kit (Origene, Rockville, MD) for mouse *Kif26b* (SKU KN308785), lipofectamine 3000, P3000 reagent and serum-free media were used for transfection of the guide RNA (ThermoFisher, Waltham, MA). 1µg of guide RNA was transfected with 1μg of donor DNA plasmid. This targets the downstream part of *Kif26b* exon 1 and the start of exon 2, replacing the coding regions with a GFP-Puromycin cassette, resulting in a gene knockout. Cells were seeded 24 hours before transfection at a density to achieve 50 – 70 % confluency on the day of transfection. Quantities of Lipofectamine 3000, P3000 and serum-free media were calculated according to the manufacturer’s instructions. After selection with 2µg/ml puromycin (ThermoFisher), resistant colonies were isolated and the knockout of *Kif26b* transcript was verified using RT-qPCR (Figure 4B) and western blotting (Figure 4C).

### Osteogenic differentiation

MSCs and hMADS were seeded for 24 hours in growth media, then 300ng/ml human recombinant BMP2 (GenScript, Wanchai, Hong Kong) was added for 48 hours. The media was replaced with the osteogenic media on day 0 of the differentiation. Osteogenic media was made as to the growth media, except for the addition of 300ng/ml BMP2, 10mM β-Glycerophosphate, 10nM Dexamethasone and 50μg/ml Ascorbic Acid (Sigma-Aldrich).

C2C12 WT and *Kif26b*^*CRISPR*^ cells were cultured in growth medium for 24 hours after seeding onto culture plates. Subsequently, growth medium ±BMP was added for 48 hours as for the MSC cultures, at which point the cells were a confluent monolayer. After confluency, the media was changed to one of the three differentiation media described above: osteogenic+BMP2, osteogenic alone, or low serum media. WT and *Kif26b*^*CRISPR*^ cells were cultured in identical conditions for the differentiation phase up to day 24 (Figure 4D). 2µg/ml puromycin selection of the *Kif26b*^*CRISPR*^ cells was only used during the growth phase. Differentiation media was made with low FBS (0.5%) and renewed every 2 days.

### RNA isolation and RT-qPCR

Total RNA was isolated using the Promega ReliaPrep™ RNA Cell, Tissue Miniprep System (Promega, Madison, WI) and RNeasy UCP Micro Kit (Qiagen) according to the manufacturer’s instructions. Reverse transcription of 400ng of RNA into cDNA was completed using the iScript™ cDNA Synthesis Kit (Bio-Rad, Hercules, CA) and the Veriti 96-well thermal cycler according to the manufacturer’s instructions (Applied Biosystems, Waltham, MA). For RT-qPCR, 2ng of cDNA was loaded per well. qPCR samples were run on the C1000 Touch™ Thermal Cycler (Bio-Rad) in 384-well plates. Triplicate technical repeats were conducted for each assay and normalised to a *β-Actin* housekeeping control in murine and *GAPDH* in human. Primers for SYBR green qPCR (Sigma-Aldrich) were designed using Primer-BLAST (NCBI: (www.ncbi.nlm.nih.gov/tools/primer-blast, Supplementary Table 10). All primers were screened to avoid self-complementarity with Oligo Calc: Oligonucleotide Properties Calculator (http://biotools.nubic.northwestern.edu/OligoCalc.html). PrecisionPLUS SYBR-Green master mix and TaqMan master mix (Primer design, Southampton, UK) were used with SYBR primers. TaqMan gene expression probes (Invitrogen, Waltham, MA) were used for mouse *Kif26b* and *β-Actin*. Ct values were presented as fold change in expression compared to day 0 and after normalisation to the housekeeping control (2^− ∆∆CT^).

### Protein Isolation and Western Blotting

Cells were lysed in 0.1% RIPA-SDS lysis buffer with protease inhibitors after three washes with PBS. For detecting phospho-proteins, phosphatase inhibitors (Sigma-Alrich) were added to the lysis buffer. Lysates were sonicated before centrifuging at 10,000 × g for 10 minutes. Protein concentration of the lysates was measured using the bicinchoninic acid assay kit (Thermo Scientific). 10 or 15μg of protein was loaded per well on Tris-Acetate 3-8% gels with Tris-Acetate running buffer (NuPAGE, ThermoFisher) for detecting KIF26B protein or on 4-12% NuPAGE Bis-Tris Gel with NuPAGE MES SDS Running Buffer (NuPAGE) for detecting phospho-proteins. Protein was transferred onto PVDF membranes with NuPAGE transfer buffer (NP0006) with 10% methanol and 1% NuPAGE antioxidant (NP0005). Blocking was for one hour at room temperature with 5% milk-TBST. The membrane was incubated with the primary antibody for 1 hour at room temperature or overnight at 4°C. Rabbit anti-human KIF26B antibody was used at 1:500 (ab121952, Abcam, Cambridge, UK) in 5% milk-TBST, the immunised sequence identical in the mouse. β-Actin and Hsp90β housekeeping controls were used at 1:5000 in milk-TBST. Anti-phospho-SMAD 1/5/8 antibody (Cell Signalling Technologies, Danvers, MA), anti-phospho-P38 (Cell Signalling Technologies) and anti-phospho-ERK1/2 (Cell Signalling Technologies) were used at 1:500-1:1000 in 5% BSA-TBST. The membrane was then incubated with the secondary horseradish peroxidase antibody for one hour at room temperature. Secondary-HRP conjugated antibodies were diluted 1:5000 for detecting KIF26B and housekeeping protein, 1:2500 for detecting phospho-proteins. Immunocomplexes were visualised with ECL western blotting detection reagent (GE Life Sciences, Amersham, UK). Western bands were quantified using Image Studio Lite Ver 5.2 (Li-COR Biosciences, Lincoln, NE).

### Alizarin Red S staining

Cells were washed twice with PBS and fixed overnight at 4°C in 100% ethanol. Cells were then washed twice with PBS before the addition of 40mM Alizarin Red S (Sigma-Aldrich), pH 4.2. The wells were washed extensively with 95% ethanol until all unbound stain was removed, the same number of washes was used for each well. Plates were air dried overnight and scanned on high-resolution flat-bed scanner at 1200dpi. ImageJ Software (NIH: http://rsb.info.nih.gov/ij/) was used to quantify the percentage mineralised area in each well. The area fraction positive for the stain was recorded, representing percentage mineralisation. Identical settings were used for all wells. The wild type wells were used as reference for setting the threshold values.

### Immunocytochemistry

Cells were cultured in 24-well plates or on chamber slides. Cells were washed twice in PBS and fixed in room temperature 4% (w/v) paraformaldehyde (PFA)-PBS for 30 minutes. Cells were washed again with PBS (three times) before blocking with 1% BSA-PBS for 45 minutes. Primary antibody was added for 1 hour at room temperature (covered to protect from light) before three washes with PBS and the addition of secondary antibody for 1 hour at room temperature (covered to protect from light). Cells were then washed again with PBS before imaging. Anti-α-tubulin mouse monoclonal antibody was used to visualise microtubules (SC32293, SantaCruz Biotechnology, Dallas, TX). NL493 green donkey anti-mouse (NL009: R&D Systems, Minneapolis, MN) secondary was used. Hoechst stain was used to stain the nuclei. Polyclonal rabbit anti-human KIF26B antibody (ab121952, Abcam) was used with anti-Rabbit NL557 Conjugated Donkey IgG secondary (NL004: R&D Systems). Secondary antibody only was used for the control conditions and settings were adjusted to remove non-specific binding. The same settings were used for all images. Anti-KIF26B was diluted 1:50 and anti-α-tubulin was diluted 1:100 for immunocytochemistry (ICC).

### Immunohistochemistry

Achilles tenotomy in the rat is an established model of heterotopic ossification that closely recapitulates heterotopic ossification associated with joint replacement.^36,37,66^ Formalin-fixed, paraffin embedded rat lower limbs with previously-healed blade-induced Achilles tenotomy (a kind gift from L Grover, Birmingham, UK) were dewaxed and rehydrated to water through a graded alcohol series. Slides were then washed in PBS, endogenous peroxidase activity quenched with 3% hydrogen peroxide for 10 mins, washed in PBS and then incubated in 2.5% horse blocking serum (ImmPRESSpress anti-rabbit IgG reagent Kit, Vector Laboratories, Burlingame, CA) for 1 hour at room temperature. Sections were then incubated with either a rabbit polyclonal anti-human KIF26B antibody (17422-1-AP, Proteintech, Rosemont, IL) or a rabbit polyclonal IgG isotype control antibody (ab37415, Abcam) for 90 mins at room temperature. Slides were then rinsed in PBS, incubated with the ImmPRESS™ Reagent for 30mins at room temperature before addition of ImmPACT DAB chromogen (SK-4105, Vector Laboratories). Sections were then counterstained with Hematoxylin, dehydrated through a graded alcohol series and mounted with DPX. All slides were scanned using a Panoramic 250 Flash III digital slide scanner (3D HISTECH, Budapest, Hungary) and images processed using QPath.

### Statistical analysis

Data are presented as mean ± SEM. Student’s t-test or, one-way or two-way ANOVA was used, with various post-hoc tests as indicated in the figure legends. All analyses are 2-tailed and statistical significance is represented as P<0.05. *P<0.05, **P<0.01, ***P<0.001, ****P<0.0001. GraphPad Prism 7 (GraphPad software) was used to present and analyse quantitative data.

## Data availability

The RNA sequencing data examined in primary human chondrocytes is available through the European Genome-phenome Archive accession numbers: EGAS00001002255, EGAD00001003355, EGAD00001003354, EGAD00001001331. The genotype data is available through accession numbers EGAD00010001746, EGAD00010001285, EGAD00010001292, EGAD00010000722. The RNASeq data for the *in silico* analysis of human MSCs in the GEO dataset is available at https://www.omicsdi.org/dataset/geo/GSE89179.

