## Supplementary material for "Genome-wide association and functional analyses identify CASC20 and KIF26B as target loci in heterotopic ossification": SF1

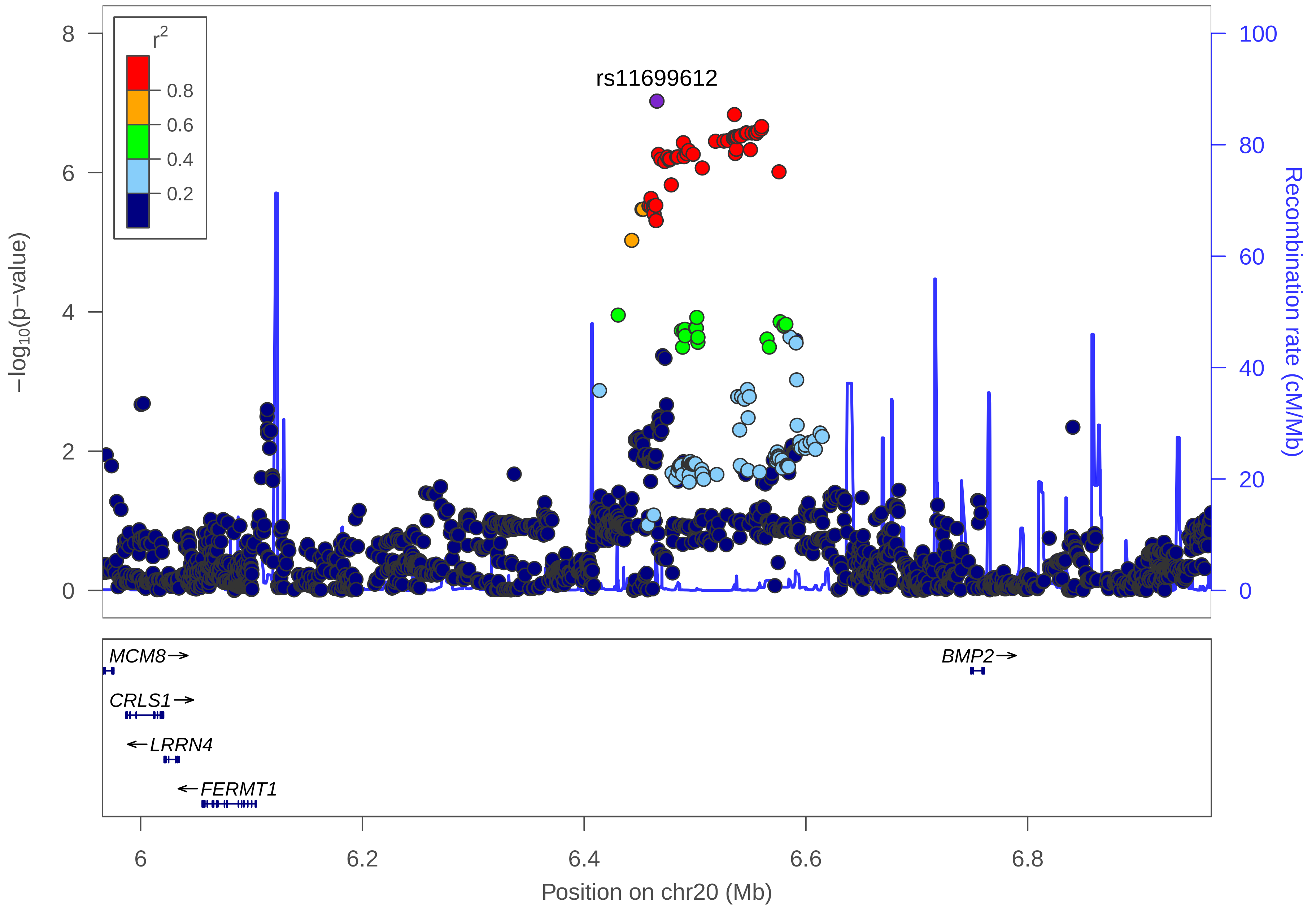

r2
0.8
0.6
0.4
0.2
rs11699612
8
6
4
2
0
−log10(p−value)
100
80
60
40
20
0
Recombination rate (cM/Mb)
MCM8→
CRLS1→
←LRRN4
←FERMT1
BMP2→
6
6.2
6.4
6.6
6.8
Position on chr20 (Mb)

| | |
|---|---|
| date: | Fri Jan 8 10:49:53 2016 |
| build: | hg19 |
| display range: | chr20:5965633−6965633 [5965633−6965633] |
| hilite range: | 0 − 0 [ 0 − 0 ] |
| reference SNP: | chr20:6465633 |
| number of SNPs plotted: | 2015 |
| requentist_add_age_sex_score_pvalue: | 9.39E−8 [chr20:6465633] |
| requentist_add_age_sex_score_pvalue: | 9.99E−1 [chr20:6084444] |

annotation key

- framestop
- splice
- nonsyn
- coding
- utr
- tfbscons
- mcs44placental
- no annotation
- none

Make more plots at http://csg.sph.umich.edu/locuszoom/
