## Supplementary material for "Genome-wide association and functional analyses identify CASC20 and KIF26B as target loci in heterotopic ossification": SF2

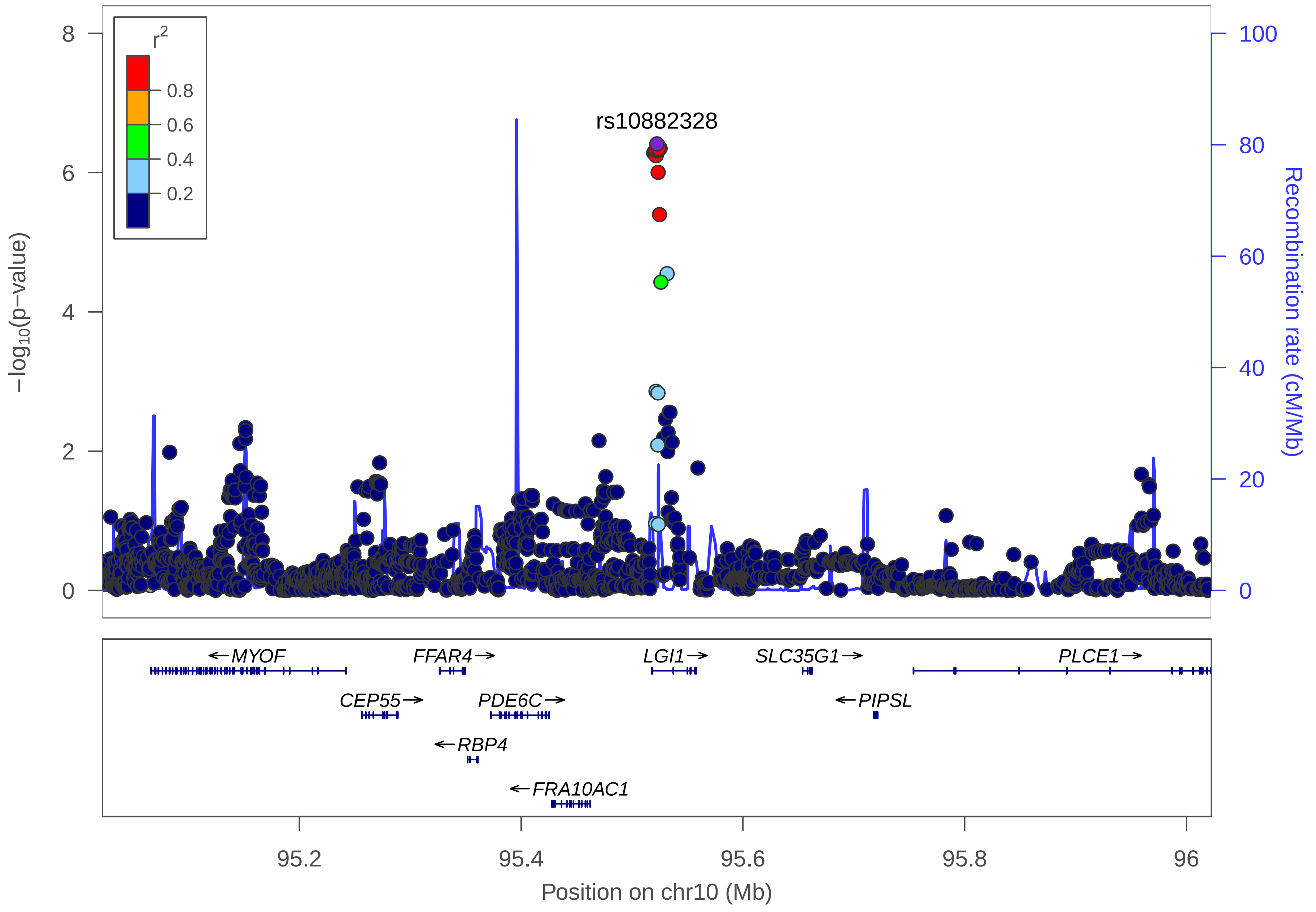

r2
0.8
0.6
0.4
0.2
rs10882328
−log10(p−value)
Recombination rate (cM/Mb)
0
2
4
6
8
0
20
40
60
80
100
MYOF
FFAR4
LGI1
SLC35G1
PLCE1
CEP55
PDE6C
PIPSL
RBP4
FRA10AC1
95.2
95.4
95.6
95.8
96
Position on chr10 (Mb)

date: Fri Jan 8 10:52:41 2016

build: hg19

display range: chr10:95022456−96022456 [95022456−96022456]

hilite range: 0 − 0 [ 0 − 0 ]

reference SNP: chr10:95522456

number of SNPs plotted: 1995

requentist_add_age_sex_score_pvalue: 3.87E−7 [chr10:95522456]

requentist_add_age_sex_score_pvalue: 9.99E−1 [chr10:95266411]

annotation key

framestop
splice
nonsyn
coding
utr
tfbscons
mcs44placental
no annotation
none

Make more plots at http://csg.sph.umich.edu/locuszoom/
