## Supplementary material for "Genome-wide association and functional analyses identify CASC20 and KIF26B as target loci in heterotopic ossification": SF3

**Supplementary Figure 3: Power to detect genetic associations.**

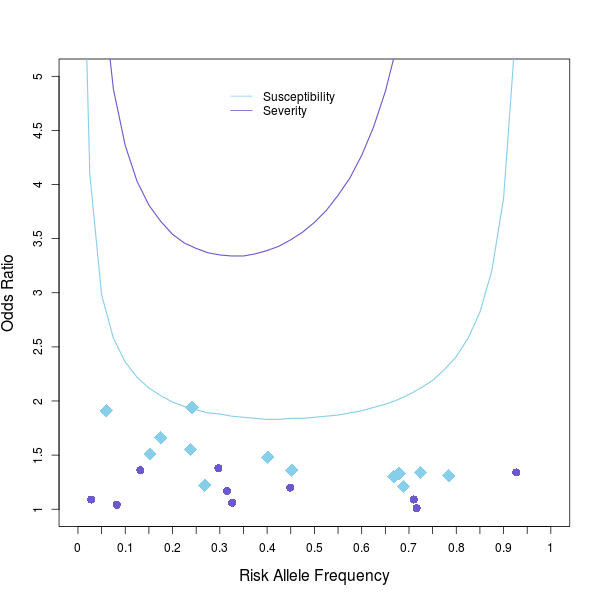

OR values are shown as a function of risk allele frequency. Light blue diamonds and purple circles denote the variants of the susceptibility and severity meta-analysis, respectively. The curves indicate 80% power at the genome-wide-significance threshold of P < 5.0 × 10−8.
