## Supplementary material for "Genome-wide association and functional analyses identify CASC20 and KIF26B as target loci in heterotopic ossification": ST8

**Supplementary Table 8. Characteristics of patients included in genome-wide association analyses.** Numerical data are mean ± standard deviation. Analyses are *Student’s t-test or **chi-squared test.

| **Discovery cohort** | | | |
| --- | --- | --- | --- |
| Characteristics | Control Group (n=481) | HO Group (n=410) | P value |
| Age at surgery (years) | 63.7±8.9 | 63.4±8.9 | 0.7* |
| Sex (female/male) | 296/185 | 194/216 | <0.0001** |
| Brooker Grade (0, 1, 2, 3, 4) | 481, 0, 0, 0, 0 | 0, 201, 133, 69, 1 | NA |
| **Replication cohort** | | | |
| Characteristics | Control group (n=212) | HO group (n=207) | P value |
| Age at surgery (years) | 67.3±10.4 | 69.4±9.5 | 0.057* |
| Sex (female/male) | 141/64 | 78/120 | <0.0001** |
| Brooker grade (0, 1, 2, 3, 4) | 205, 0, 0, 0, 0 | 0, 67, 65, 49, 17 | NA |
