## Supplementary material for "Genome-wide association and functional analyses identify CASC20 and KIF26B as target loci in heterotopic ossification": ST10

*Supplementary Table 8.* Human and Mouse qPCR primer sequences.

| Gene | Human qPCR Primer Sequence (5’ -> 3’) |
| --- | --- |
| *GAPDH* | FW ATTGCCCTCAACGACCACTTT  REV CCCTGTTGCTGTAGCCAAATTC |
| *CASC20* | FW TCATATGGATTTCAAGCTGGGT  REV TCCCAGTCTTCTGCATCACTTC |
| *KIF26B* | FW GTGTTCTGTTTCGGCCACG  REV CCCAGGTTCTGCATGGAATCA |
| *RUNX2* | FW GGTTAATCTCCGCAGGTCACT  REV CACTGTGCTGAAGAGGCT |
| *OSX* | FW CCACCTACCCATCTGACTTTG  REV CCACTATTTCCCACTGCCTT |

| Gene | Mouse qPCR Primer Sequence (5’ -> 3’) |
| --- | --- |
| *Alp* | FW CCCCGGGGCAACTCCATCTT  REV TAGCCAGGCCCGTTACCATA |
| *β-Actin* | FW GGGACCTGACAGACTACCTCATG  REV GTCACGCACGATTTCCCTCTCAGC |
| *Bglap* | FW CCAAGCAGGAGGGCAATAAGGTA  REV GGATCTGGGCTGGGGACTGAG |
| *Col1a1* | FW CTGGCAACAAAGGAGACACTGG  REV GGGCCTGGGGGACCTTGAACTC |
| *Osterix* | FW TCACACCCGGGAGAAGAAGTT  REV CCCGTGGGTGCGCTGATGTT |
| *Runx2* | FW ACTGGCGGTGCAACAAGA  REV GACGGTAACCACAGTCCCATC |
